# Novel phylogeny of angiosperms inferred from whole-genome microsynteny analysis

**DOI:** 10.1101/2020.01.15.908376

**Authors:** Tao Zhao, Jiayu Xue, Shu-min Kao, Zhen Li, Arthur Zwaenepoel, M. Eric Schranz, Yves Van de Peer

## Abstract

Despite the wealth of genomic and transcriptomic data of pivotal angiosperm species, the phylogenetic relationships of flowering plants are still not fully resolved. Microsynteny, or the conservation of relative gene order, has been recognized as a valuable and alternative phylogenetic character to sequence-based characters (nucleotides or amino acids). Here, we present a novel approach for phylogenetic tree reconstruction based on genome-wide synteny network data. We generated and analyzed synteny networks from 123 species from 52 families across 31 orders of flowering plants, including several lineages for which phylogenetic relationships are ambiguous. We obtained a stable and highly resolved phylogeny that is largely congruent with sequence-based phylogenies. However, our results unveiled several novel relationships for some key clades, such as magnoliids sister to monocots, Vitales as sister to core-eudicots, and Saxifragales sister to Santalales, in turn both sister to Caryophyllales. Our results highlight that phylogenies based on genome structure and organization are complementary to sequence-based phylogenies and provide alternative hypotheses of angiosperm relationships to be further tested.

## Introduction

Phylogenetic trees serve many purposes and are an indispensable tool for the interpretation of evolutionary trends and trait changes. As a consequence, phylogenetics – and more recently phylogenomics – has been a very active field of research (Delsuc et al., 2005; Philippe et al., 2005; Hackett et al., 2008). Phylogenetic reconstruction is generally based on the comparison of informative homologous characters, which can be morphological data, and/or molecular (sequence) data. In particular the latter have been widely used for phylogeny reconstruction, for reasons that have been extensively discussed elsewhere (Doolittle, 1999; Delsuc et al., 2005; Ciccarelli et al., 2006).

Plant molecular phylogenetic studies, based on representative plastid, mitochondrial, or nuclear genes, have greatly advanced our knowledge and understanding of angiosperm evolution. For example, the Angiosperm Phylogeny Group (APG) provides a widely accepted classification for flowering plants (Chase et al., 2016). With the development of high-throughput sequencing technologies, molecular phylogenetics has entered a new era, characterized by the usage of large-scale genomics data, such as entire transcriptomes (One Thousand Plant Transcriptomes Initiative, 2019) or whole genome sequence data. In such a typical ‘phylogenomics’ approach, a set of genes present across the full taxon set under consideration is selected and concatenated into a supermatrix, after which a tree can be inferred with standard phylogenetic tree reconstruction methods (Baldauf et al., 2000; Ciccarelli et al., 2006). Alternatively, trees inferred for individual gene families may be combined using a so-called supertree approach. Among the latter, methods based on the multi-species coalescent (Rannala and Yang, 2003; Mirarab et al., 2016) are increasingly adopted, as supermatrix approaches may fail in the face of incomplete lineage sorting (ILS), in which case the gene trees may not reflect the species tree topology (Degnan and Rosenberg, 2006; Mendes and Hahn, 2017).

However, even based on extensive data, sequence-based phylogenies often yield conflicting topologies, because of differences in taxon sampling and/or gene marker selection (Philippe et al., 2011). Furthermore, homology or orthology assessment has proven particularly difficult for plants, due to complications associated with frequent gene duplication and loss (Rasmussen and Kellis, 2012), ancestral hybridization (Solís-Lemus et al., 2016), and lateral gene transfer (Yang et al., 2016). Organelle genes might be less prone to duplications and losses, but might introduce biases since they are generally maternally inherited (Greiner et al., 2015; Liu et al., 2019). Besides, misspecifications in nucleotide substitution or amino acid replacement models can be the cause of systematic errors (Jeffroy et al., 2006; Rodríguez-Ezpeleta et al., 2007; Philippe et al., 2011).

Therefore, alternative ways to infer phylogenetic relationships are needed. Gene order conservation, or microsynteny, has been recognized as highly informative for phylogenetic purposes (Sankoff et al., 1992). However, so far, application of gene order conservation for phylogenomic means has been limited (Rokas and Holland, 2000; Tang and Moret, 2003; Drillon et al., 2019). Recently, a novel approach in which (micro)synteny information is converted into a network data structure was developed, and which has proven to be well suited for evolutionary comparisons among many complex eukaryotic nuclear genomes (Zhao et al., 2017; Zhao and Schranz, 2019). Here, we build on this recently developed network-based microsynteny approach by integrating synteny networks and their matrix representation with maximum likelihood-based phylogenetic inference to reconstruct the phylogeny of flowering plants. To this end, we analyzed 123 flowering plant genomes, representing 52 families in 31 orders, and compared inferred phylogenies with phylogenies obtained with more traditional, sequence-based methods. While our microsynteny-based trees are surprisingly congruent with trees inferred by more classical (supermatrix and supertree) approaches, we observed notable differences with respect to the clustering of magnoliids, Vitales, Caryophyllales, Saxifragales, and Santalales. We believe our approach provides a noteworthy complement to more classical approaches of tree inference, and propose it could solve some conundrums that are difficult to solve with sequence (alignment) based methods.

## Results

After initial genome quality control (see Methods), 123 fully sequenced plant genomes were used for further analysis (Supplemental Table 1). The overall sampling covered 31 orders and 52 different families of angiosperms. The phylogenomic synteny network construction pipeline was used as previously described (see Methods and Zhao and Schranz, 2019) (Figure 1a left panel). Briefly, all-vs-all pairwise genome comparisons were conducted, followed by the detection of synteny blocks across and within all genomes. This is followed by network construction in which the nodes represent genes and edges are added between each anchor pair (homologous genes within synteny blocks).

**Figure 1.**
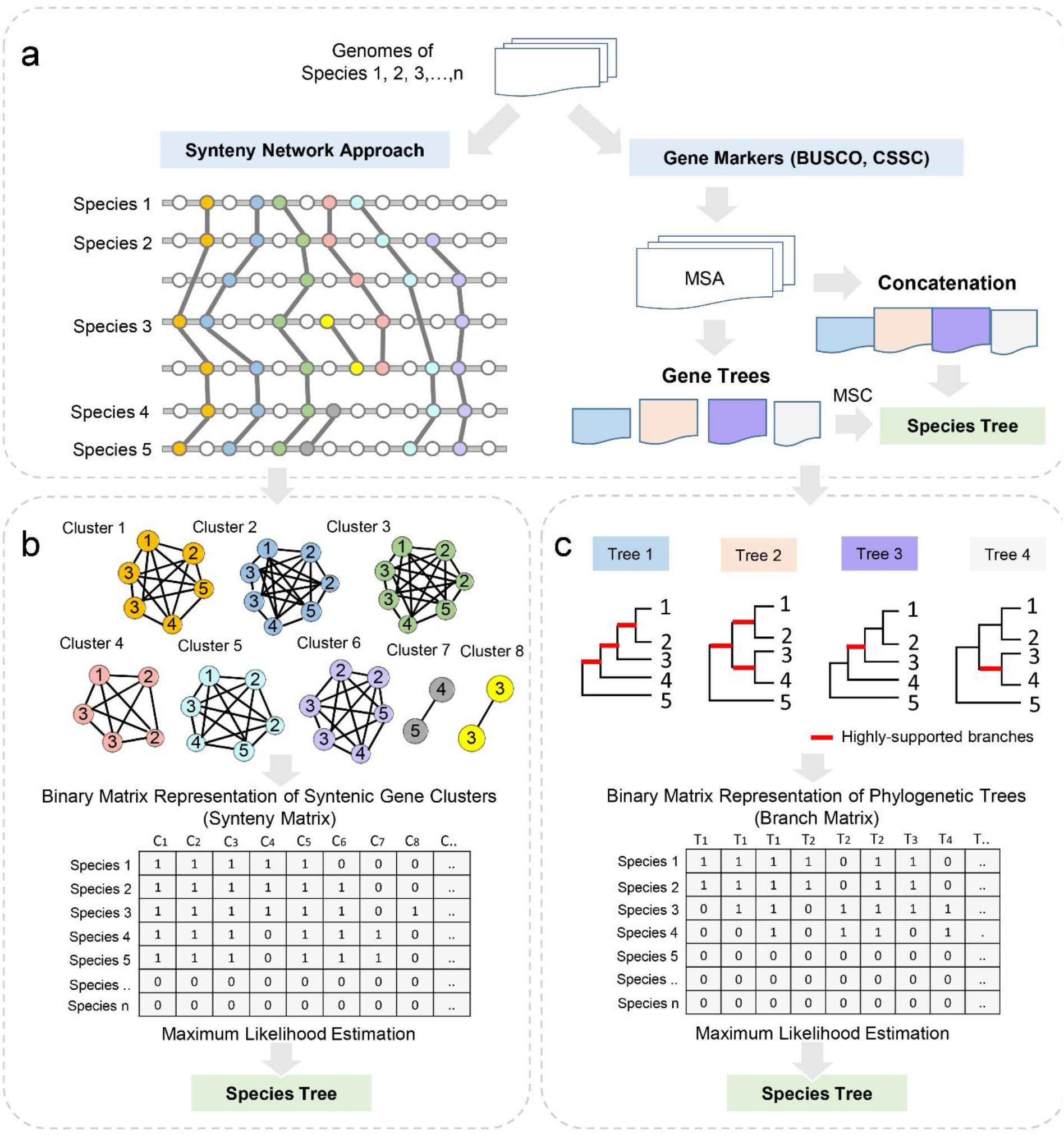
Whole-genome based species tree inference. (**a**) Whole genome annotations of all predicted genes are used for the synteny network approach (left panel) and gene marker identification (right panel). Multiple sequence alignment (MSA) was performed for each of the delineated (orthologous) gene families or clusters (orthogroups). Species trees can then be inferred by sequence concatenation (supermatrix) and supertree approaches (see text for details). The synteny network approach conducts all pairwise reciprocal genome comparisons, followed by synteny block detection. All synteny blocks constitute the synteny network database (see Methods for details). (**b**) Synteny clusters vary in size and node compositions. The phylogenomic profiling of all clusters construct a binary matrix, where rows represent species and columns as clusters. This synteny matrix is used as the input for species tree estimation by maximum likelihood (referred to as Syn-MRL). (**c**) MRL supertree method for species tree construction. Phylogenetic gene trees of individual protein alignments are used as source trees. Leaves from well-supported nodes of all trees construct a binary branch matrix, which is then used for phylogenetic analysis by maximum likelihood.

The entire synteny network summarizes information from 7,435,502 pairwise syntenic blocks or segments, and contains 3,098,333 nodes (genes) and 94,980,088 edges (sytenic connections). We clustered the entire network into 137,833 synteny clusters (exemplified in Figure 1b). Synteny clusters vary in size and number of species, which indicate whether the gene synteny is well conserved or rather lineage-/species-specific.

Phylogenomic profiles of all clusters are combined as columns of the synteny matrix, which is used for maximum likelihood (ML) tree inference (see Methods, Figure 1b). Thus, the dimension of the input synteny matrix is 123 × 137,833.

For the gene sequence-based approaches, we used BUSCO genes (Benchmarking Universal Single-Copy Orthologs) and CSSC genes (Conserved Single-copy Synteny Clusters, see Methods) independently for both the supermatrix (concatenated sequence alignments) and supertree methods (Figure 1a right panel). For the supermatrix approach, BUSCO analysis (v3.0, embryophyta_odb9, with 1440 profiles) identified a total of 1438 conserved single-copy genes from the 123 angiosperm genome sequences, compared to 883 identified as CSSC. The length of the concatenated gene sequence alignments of BUSCO and CSSC genes were 591,196 and 341,431 amino acids, respectively. For the supertree approach, we used variants of the matrix representation with likelihood (MRL) analysis (see Methods) based on a set of input trees generated from BUSCO and CSSC phylogenetic gene markers (Figure 1c). Each column (matrix element) in the matrix represents a well-supported bipartition (bootstrap support (BS) ≥ 85%) (Figure 1c). The dimensions of the matrices are 123 × 139,538 for the BUSCO gene trees, and 123 × 102,617 for the CSSC gene trees. Besides the MRL supertree, we also used ASTRAL, which is statistically consistent under the multi-species coalescent model, to estimate species trees from BUSCO and CSSC gene trees, respectively.

### Synteny versus sequence-alignment based phylogenetic trees of angiosperms

We obtained seven phylogenetic trees based on the 123 species dataset: one tree inferred from synteny data (further referred to as the ‘synteny tree’), and two times three trees based on sequence-alignments, respectively (referred to as the concatenation-BUSCO tree and the concatenation-CSSC tree, the MRL-BUSCO tree and the MRL-CSSC tree, and the ASTRAL-BUSCO tree and ASTRAL-CSSC tree (Supplemental Figures S1-S7)). For all trees, *Amborella* was used as the outgroup (One Thousand Plant Transcriptomes Initiative, 2019). Overall, the phylogenetic trees are highly resolved, well supported, and highly congruent with each other. Since the six sequence-alignment based trees are highly similar, and only differ by a few bootstrap support values and a few minor sister-group relationships, we further use the concatenation-BUSCO tree (further referred to as the ‘BUSCO tree’) as our representative of the sequence-alignment based trees and for comparison with the single ‘synteny tree’ (Figure 2).

**Figure 2.**
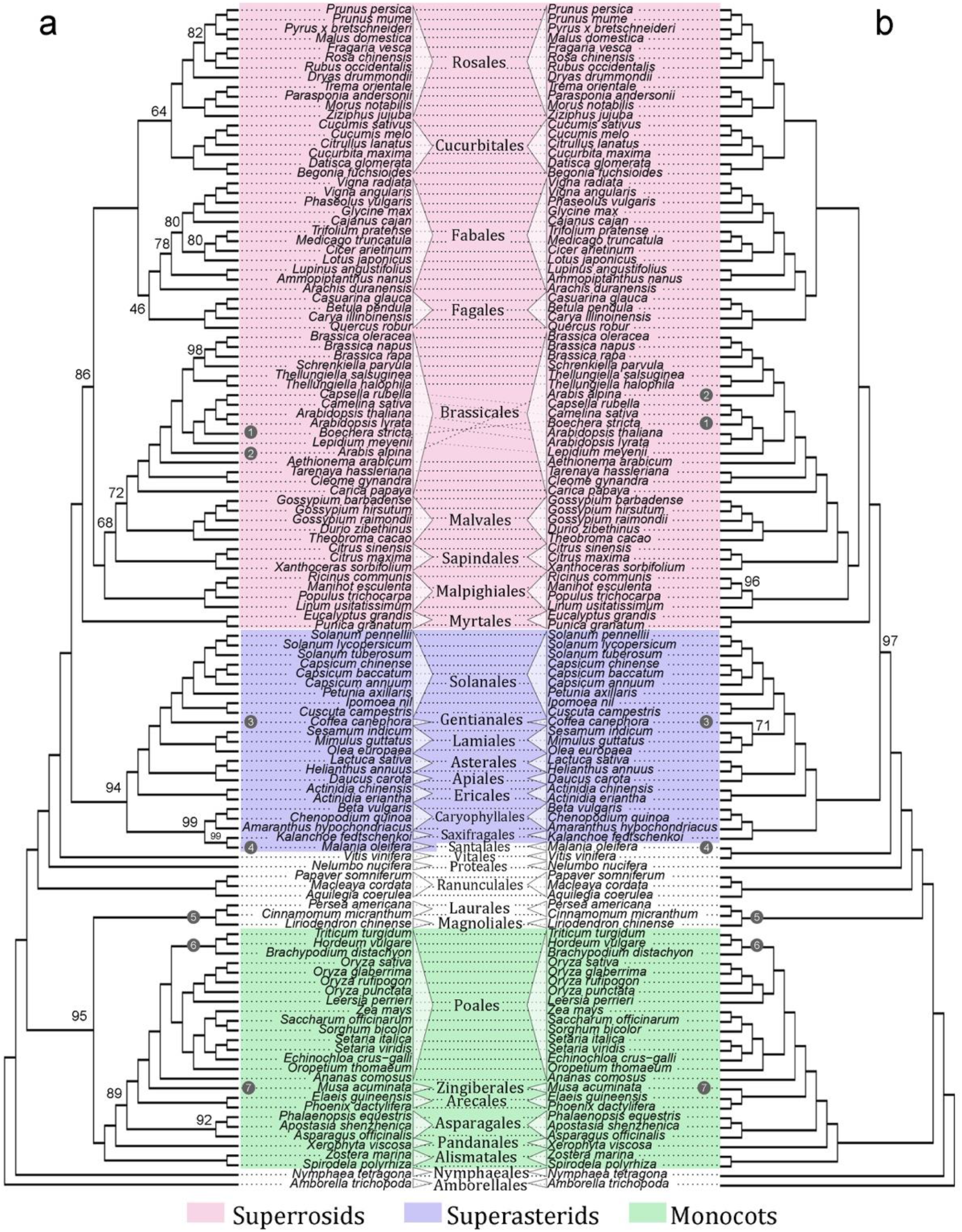
Best maximum likelihood (ML) tree for 123 fully-sequenced flowering plant genomes. (**a**) Best ML tree based on the synteny matrix. (**b**) Best ML tree based on the concatenation of protein alignments of BUSCO genes. Both trees are rooted by *Amborella*, and three main clades, i.e. superrosids, superasterids, and monocots are shaded in light-red, light-purple, and light-green, respectively. Seven differences are indicated by indexed black dots. Branches are not drawn to scale. Ultrafast bootstrapping values (see text for details) were marked for nodes with less than 100% support.

Despite the fact that both trees (synteny tree and BUSCO tree) are highly similar, there are three significant differences between the two trees. Firstly, in the synteny tree magnoliids (including Laurales and Magnoliales) form a sister group to monocots (BS = 95%), with both then sister to the eudicots (BS = 100%) (Figure 2a), while in the BUSCO tree, magnoliids are sister to the eudicots and both are then sister to the monocots (Figure 2b). Secondly, within the Poales synteny suggests that Ehrhartoideae (including the genera *Oryza* and *Leersia*), which is a part of BEP clade (Bambusoideae, Ehrhartoideae (formerly Oryzoideae) and Pooideae) (Grass Phylogeny Working Group II, 2012), is sister to the PACMAD clade (including sugarcane, maize, and sorghum, etc.) (BS = 100%), in turn sister to the Pooideae (including wheat, barely, and *Brachypodium*), which also belong to the BEP clade (BS = 100%) (Figure 2a). On the contrary, the BUSCO tree supports the original BEP clade, including Pooideae and Ehrhartoideae, as sister to the PACMAD clade (BS = 100%) (Figure 2b). Thirdly, our synteny tree supports Santalales as sister to Saxifragales (Figure 2a), while in the BUSCO tree, Santalales are sister to Vitales (Figure 2b). Additional more subtle differences also were found including the position of *Musa* (Zingiberales), *Arabis* (Brassicaceae), *Boechera* (Brassicaceae), *Coffea* (Gentianales), and *Malania* (Santalales) (Figure 2). In particular, synteny supports Zingiberales + Poales as sister to Arecales (Figure 2a), whereas BUSCO supports Zingiberales + Arecales as sister to Poales (Figure 2b). Likewise, synteny supports Gentianales + Solanales as sister to Lamiales (Figure 2a), whereas BUSCO supports Gentianales + Lamiales as sister to Solanales (Figure 2b).

### Comparison with APG IV

We further summarized the differences between the synteny-based and alignment-based phylogenies (represented by concatenation-BUSCO) at the order level and compared each to the current APG phylogeny (version IV) (Chase et al., 2016) (Figures 3a-c). Despite the abovementioned differences, the synteny and BUSCO trees agree on a number of branching orders that are different than those of the current APG tree. Both of our trees strongly favor Vitales sister to the rest of the core eudicots, or as another clade of early-diverging eudicots, branching off after Proteales (*Nelumbo*) (both BS = 100%). In the synteny tree, Santalales is sister to Saxifragales, and with both sister to Caryophyllales, which in turn sister to all other asterids. The BUSCO tree supports the same topology, except Santalales is clustered with Vitales (Figures 2 and 3). Both trees positioned Myrtales (*Eucalyptus* and *Punica*) as basal rosids, and Malpighiales as basal Malvids. For orders within Fabids, both trees support a clustered relationship of ((Fagales, Fabales), (Rosales, Cucurbitales)). All these results differ from the current APG version IV tree.

**Figure 3.**
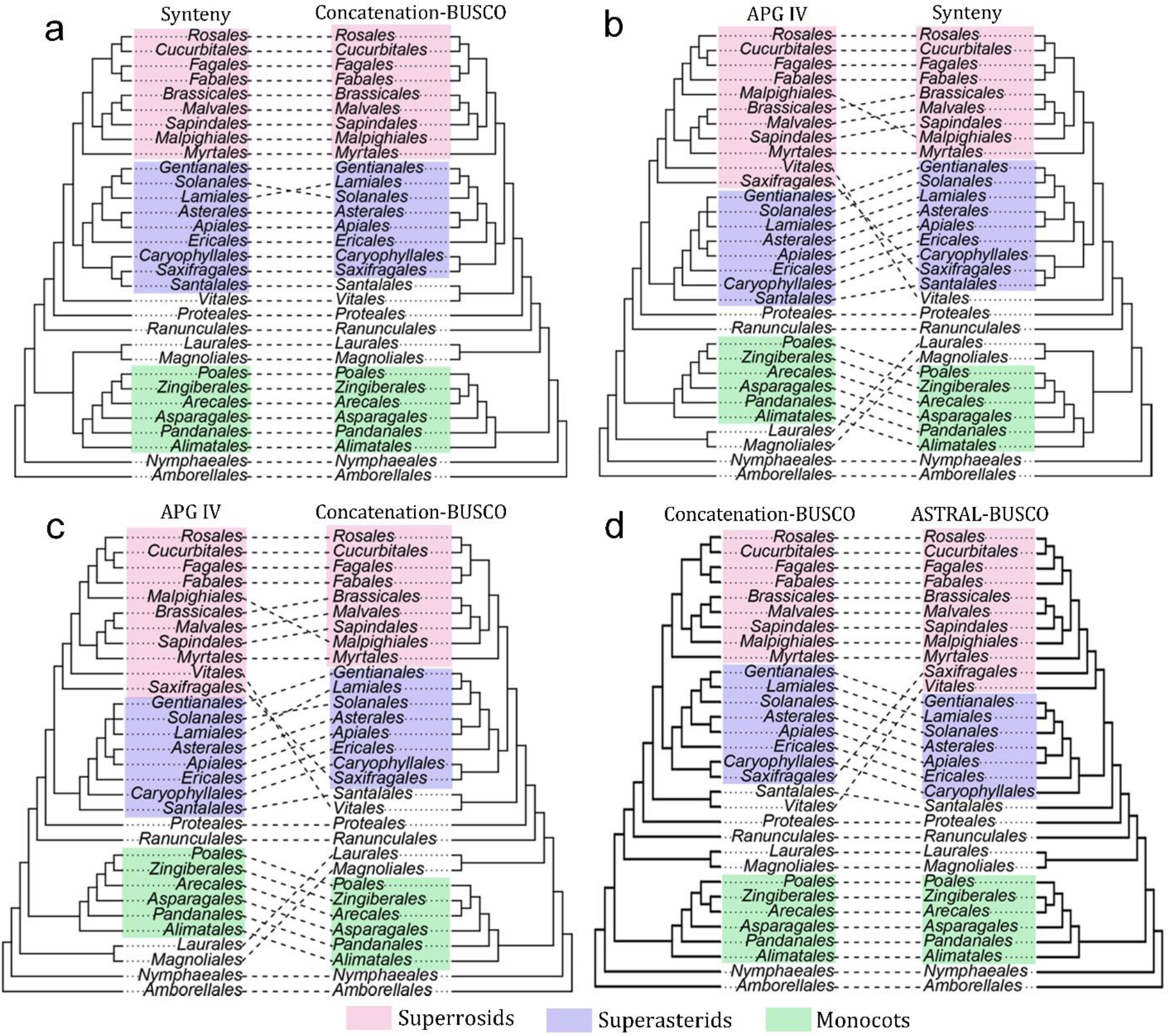
Comparison of phylogenetic relationships of 31 plant orders based on whole-genome synteny, gene markers, multi-species coalescent, and the Angiosperm Phylogeny Group (APG IV, (Chase et al., 2016)). (**a**) Comparison of the synteny and BUSCO tree. (**b**) Comparison of the APG IV and synteny tree. (**c**) Comparison of the APG IV and BUSCO tree. (**d**) Comparison of the BUSCO and ASTRAL-BUSCO tree.

ASTRAL trees are highly consistent with the BUSCO tree and other alignment-based phylogenies (from supermatrix and MRL), but positioned Saxifragales as an early-branching Rosid (Figure 3d, Supplemental Figures S6 and 7). Vitales varied in its placement between ASTRAL-BUSCO and ASTRAL-CSSC trees, in which the former supports it as early-diverging within Rosids (Supplemental Figure S6), whereas the latter supports it as early-diverging within eudicots (clustered with Santalales) (Supplemental Figure S7). Moreover, ASTRAL trees support a relationship of (Fabales, (Fagales, (Rosales, Cucurbitales))), but with lower support values (Figure 3d, Supplemental Figures S6 and 7).

We used the ‘approximately unbiased’ (AU) test to evaluate featured alternative phylogenies from the synteny, BUSCO, and APG trees. Tested hypotheses included the seven observed differences between the synteny and BUSCO tree in Figure 2 (we describe one alternative (Difference #5 from Figure 2) in Figure 4a, for the other alternatives please refer to the differences marked in Figure 2), and six differences compared to the APG tree (Figures 4b–4g). Results show that the best ML synteny tree (depicted in Figures 2a and 3a and b) was found to be significantly better than 12 out of 13 alternative topologies and all alternative hypotheses could be rejected with P < 0.05, except the scenario where magnoliids are sister to eudicots (P = 0.146, Figure 4a).

**Figure 4.**
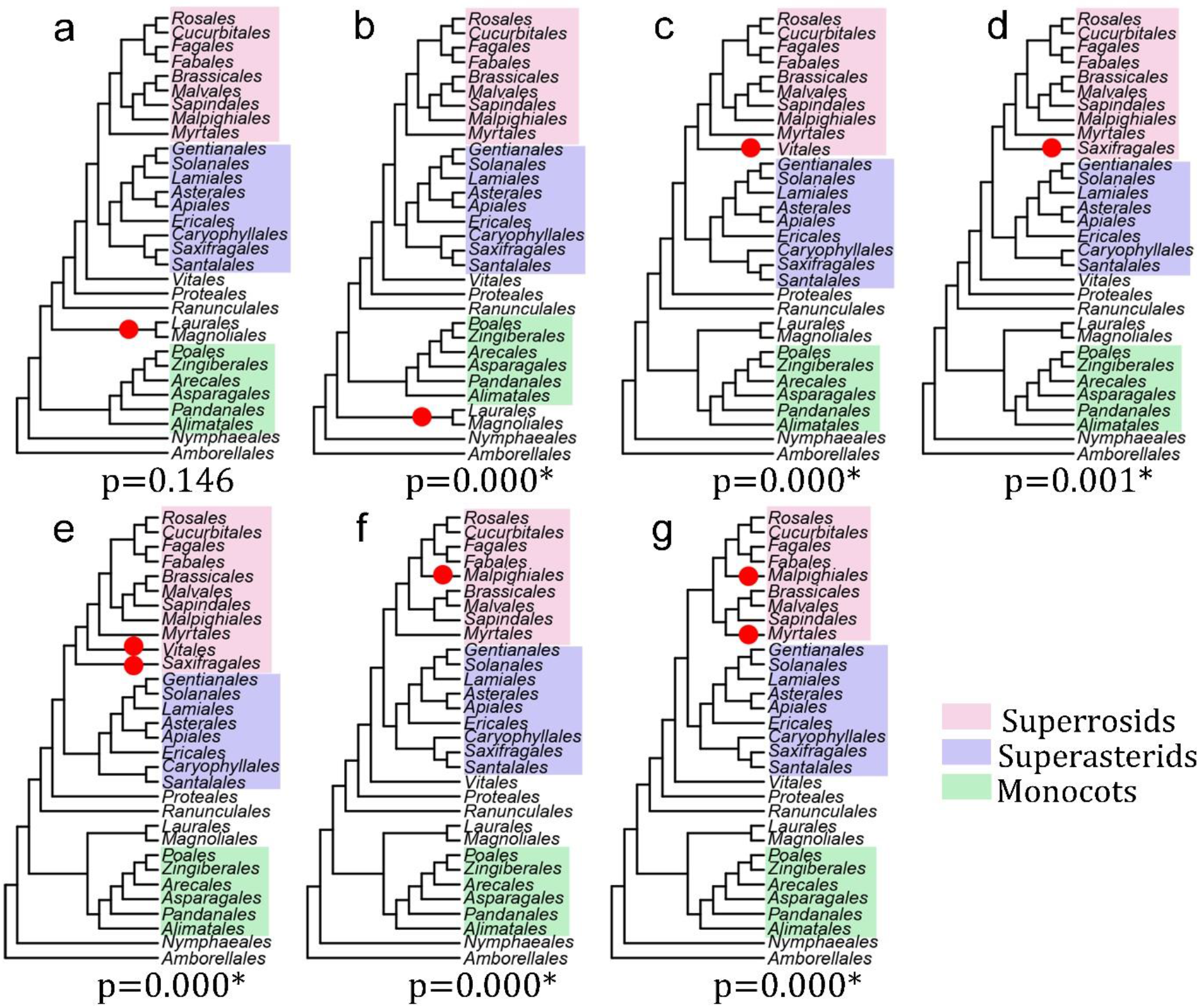
Alternative topologies used for approximate unbiased (AU) test. Alternatively tested topologies for placement of (**a**) magnoliids as sister to eudicots, (**b**) magnoliids as sister to both monocots and eudicots, (**c**) Vitales as basal rosids, (**d**) Saxifragales as basal rosids, (**e**) Vitales and Saxifragales as basal rosids, (**f**) Malpighiales as basal Fabids, and (**g**) Malpighiales as basal Fabids plus Myrtales as basal Malvids.

## Discussion

### Matrix representation-based tree reconstruction for phylogenomic microsynteny properties

The notion that the genomic architecture of extant plants is highly diversified and the result of considerable genomic rearrangement is well accepted (Murat et al., 2012; Zhao and Schranz, 2019). Plant genomes are derived and modified by a complex interplay of whole-genome duplication (Van de Peer et al., 2017; One Thousand Plant Transcriptomes Initiative, 2019), hybridization (Payseur and Rieseberg, 2016), transposable element activity (Fedoroff, 2000), and gene gain and loss (Freeling, 2009). All these phenomena present challenges for phylogenetic inference, yet at the same time they create opportunities, as these processes result in useful information for phylogenetic inference. Previous endeavors for phylogeny reconstruction using gene order information have mostly analyzed chromosomal breakpoints. For example, the Double-Cut-and-Join (DCJ) model estimates distances between chromosomes based on inversions, transpositions, fusions, and fission events, which led to genomic rearrangements (Yancopoulos et al., 2005). However, such approaches generally suffer from combinatorial explosion (Fertin, 2009) and are computationally very demanding, making them applicable only to simpler organelle or bacterial genomes. Recently (Drillon et al., 2019) developed a bottom-up pairwise (partial splits) distance-based tree reconstruction approach that starts with the identification of breakpoints between synteny blocks, followed by the identification of partial splits (Semple and Steel, 2000). However, the method was applied to ‘only’ 13 vertebrate genomes (which were phylogenetically far-apart) and 21 ‘small’ yeast genomes. Thus its performance and accuracy to more and more complex (such as plant) genomes remains to be evaluated (Drillon et al., 2019). In this study, our alternative approach bypasses these combinatorial issues by extracting genome structure information across genomes in a network. Taking advantage of phylogenomic synteny network analysis, complete gene order relatedness information for all annotated genes in (a large number of) genomes is captured and simplified into synteny clusters (Zhao and Schranz, 2019). Each cluster, no matter how conserved, family-or species-specific, contains some phylogenetic signal and can be used for tree inference.

We exploited the total information residing in all synteny clusters by encoding their phylogenomic synteny pattern profiles into a large binary matrix (synteny matrix) (Figure 1b). Standard ML-based tree inference was then applied to reconstruct a species tree (Figure 1b). The process is similar to the MRL (Matrix Representation with Likelihood) supertree method (which uses the same data matrix as MRP (Matrix Representation with Parsimony), but with ML-based inference, yielding higher accuracy (Nguyen et al., 2012; Mirarab et al., 2016)), except that MRL is based on a set of input trees, in contrast with our synteny matrix based on synteny clusters. To avoid misunderstanding, in this study we confine the usage of ‘MRL’ to the supertree method. Alternatively, we could refer to our ‘supercluster’ approach as Syn-MRL (Synteny Matrix Representation with Likelihood).

In order to assess the congruence and discordance of trees obtained by our synteny-based Syn-MRL approach and more traditional multiple sequence alignment-based approaches, we also built trees based on gene marker information. Since BUSCO gene sets are widely adopted in benchmarking genome assembly and annotation quality, we used this gene set for phylogeny reconstruction. Besides BUSCO, we also used genes from conserved single-copy synteny clusters (CSSC) for comparison. Interestingly, concatenated sequence alignments for each set of genes (1438 and 883 genes, respectively) led to exactly the same topologies (Supplemental Figures S2 and S3). For the MRL supertree approach, we added only highly-supported bipartitions of each tree to the data matrix, since in reality gene trees can rarely be fully resolved. The 85% bootstrap cutoff (ultrafast bootstrap approximation (Minh et al., 2013)) is an empirical cutoff to accommodate strictness, but also allow for signals with minor uncertainty to be included for statistical testing. Again, the resulting tree from the MRL approach is highly congruent with the concatenation-derived tree.

For the MRL and Syn-MRL approaches, we use the Mk model, which is a Jukes-Cantor type model for discrete morphological data (Lewis, 2001). Each column of the data matrix is regarded as an independent character. The two-state binary encoding for each column represent the two groups of related species regarding that specific character. There is no further ordering or special weighing for the elements in the data matrix. The reliability of the signals (columns in the matrix) used in Syn-MRL (supercluster) and MRL (supertree) depended on the network clustering and the robustness of bipartitions, respectively (Figures 1b and 1c). The Mk model may arouse some concern regarding the symmetric two-state model (i.e., there is an equal probability of changing from state 0 to state 1 and from state 1 to state 0). In general, time-reversible Markov models of evolution may not be ideal for the Syn-MRL approach. However, a first and important observation was that these models do seem to result in very reasonable well-supported phylogenies, in a similar vein as the MRL approach (Nguyen et al., 2012). We hypothesize why this is: firstly, we may reasonably suspect that for our data, state 0 or 1 has the same probability of being the ancestral or derived state, as we can hypothesize that the emergence of a new lineage-specific synteny cluster and loss of another, are due to the same processes, e.g. transposition. Secondly, although the Mk model allows numerous state changes, in practice it yields trees with identical likelihood scores than a modified Mk model where character states can change only once or not at all (Lewis, 2001). This suggests that more biologically plausible non-reversible models, where for instance the re-emergence of a synteny cluster (0→1 transition after the *initial* birth of the cluster) occurs at a different rate than the *initial* 0→1 transition, might not result in a substantial better fit. Nevertheless, we believe developing other probabilistic models for synteny cluster evolution is a fruitful avenue for further research.

### Whole-genome microsynteny data: potential solution to reconstructing true species trees under ancestral hybridization?

The notion that species boundaries can be obscured by introgressive hybridization is increasingly accepted (Dasmahapatra et al., 2012; Fontaine et al., 2015; Forsythe et al., 2018; Edelman et al., 2019). A recent study has comprehensively investigated the particular species branching order of three monophyletic groups in the Brassicaceae: *Boechera* (Clade B), *Capsella* and *Camelina* (Clade C), and *Arabidopsis* (Clade A) (Forsythe et al., 2018). Forsyth et al. (2018) explicitly proved the existence of massive nuclear introgression between Clades B and C, which largely reduced the sequence divergence between them. As a result, the majority of single-copy gene trees strongly support the branching order of A(BC), while the true branching order is supposed to be B(AC) (Forsythe et al., 2018). Their study employed multiple approaches and measurements to show that neither gene duplication and loss nor ILS, nor phylogenetic noise can adequately explain such incongruence, except introgressive hybridization (Forsythe et al., 2018).

Interestingly, our results are congruent with the findings of Forsythe et al. (2018). Alignment-based trees (supermatrix, MRL, and ASTRAL) all support A(BC) (Figure 2b, Supplemental Figures 2-7), whereas only the synteny tree clearly supports the ‘true’ species branching order B(AC) (Figure 2a). Since the focal point of synteny information is to reflect genomic structure variation instead of gene sequence changes, we have reasons to believe that genome level synteny-based phylogeny inference may intrinsically avoid the bias caused by (extensive) introgression (recent or more ancestral) for species tree reconstruction, and can reflect the ‘true’ species branching order. In any case, the whole-genome synteny approach could provide unique complements towards developing suitable deep-time hybridization models and systems (Folk et al., 2018).

### The phylogenetic position of magnoliids and beyond

The phylogenetic position of magnoliids has long been discussed (Soltis and Soltis, 2019). Therefore, we have explored the relationships of magnoliids, monocots, and eudicots in more detail. To this end, we focused on the ‘submatrix’ of synteny profiles for magnoliids and extracted 15,424 magnoliid-associated clusters (signals involving magnoliids) (Figure 5a). A clustering of the phylogenomic profiling patterns of these clusters evidently showed 1,107 clusters strongly supporting a grouping of magnoliids and monocots (Figure 5a). To validate their contribution to the final best ML tree, we removed these signals from the entire matrix and rebuilt the phylogenetic tree, after which the species tree obtained then favors magnoliids as sister to eudicots (BS = 100%) (Supplemental Figure S8). To further understand the genomic distribution of specific genes in these clusters joining magnoliids and monocots, we reorganized the cluster profiles according to the chromosome gene arrangement of a magnoliid representative (*Cinnamomum micranthum*) (Supplemental Table S2). In doing so, we observed a number of ‘signature’ blocks consisting of specific anchor pairs that are shared by monocots and magnoliids to the exclusion of eudicots (Supplemental Table S2-sheet 1, contexts with highlighted yellow rows). As an example, In Figure 5b, we highlight a synteny context of 15 genes where 8 genes (highlighted red) are only found in synteny between magnoliid and monocot genomes (Figure 5b), with close flanking genes generally conserved angiosperm-wide (highlighted blue) (Figure 5b, Supplemental Table S2-sheet2).

**Figure 5.**
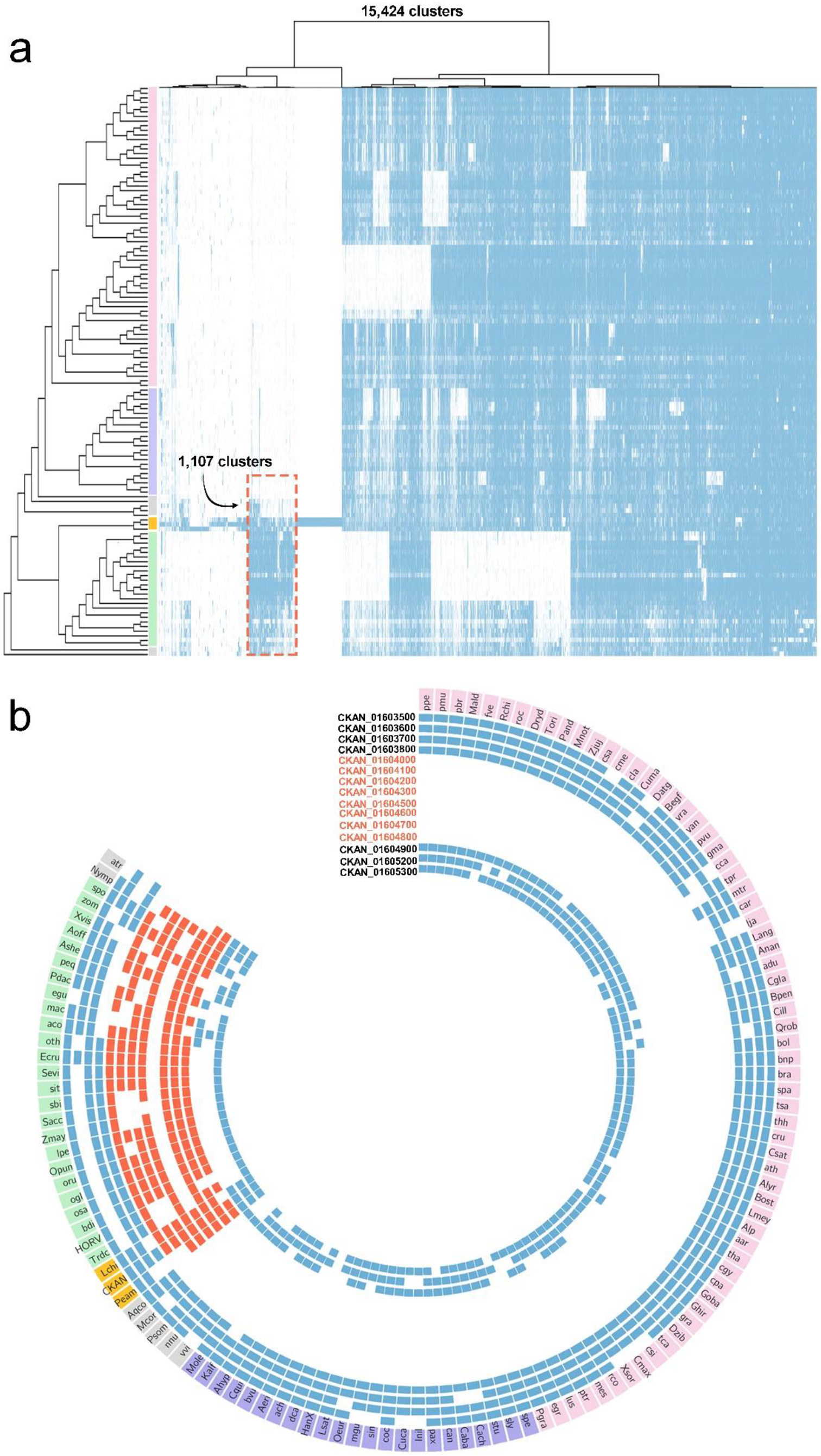
Magnoliids-associate signals and example of microsynteny. (**a**) Hierarchical clustering (ward.D) of 15,424 magnoliids-associate cluster profiles based on jaccard distance. On the far-left, the synteny species tree is displayed (same as Figure 2a). Three magnoliid genomes are highlighted in orange. 1,107 clusters represent signals supporting magnoliids close to monocots. (b) A fifteen-gene context in the genome of *Cinnamomum micranthum* (a magnoliid) shows eight neighbouring genes (highlighted in orange) only present in magnoliids and monocot genomes, while the flanking genes (colored blue) are generally conserved angiosperm-wide. Outmost are species as abbreviated names, which are of the same order and color as in (**a**). See text for details.

Apart from the contentious phylogenetic relationship of magnoliids, also the evolutionary relationships at the base of the core eudicots, such as Vitales, Saxifragales, Santalales, and Caryophyllales, has proven elusive (Zeng et al., 2017; One Thousand Plant Transcriptomes Initiative, 2019). Our ASTRAL trees showed short internal branches for these lineages, and placements of Vitales varied in the two ASTRAL trees (Supplemental Figures 6 and 7). These results are consistent with the findings of the recent 1KP study, in which taxa in Vitales (*Tetrastigma obtectum, Tetrastigma voinierianum, Vitis vinifera*, and *Cissus quadrangularis*) were recongized as ‘rogue’, since they occupied different phylogenetic positions in different analyses (ASTRAL coalescent, supermatrix, and plastid genome) (One Thousand Plant Transcriptomes Initiative, 2019). Our synteny-based phylogenies strongly support Vitales as another clade of early-diverging eudicots branching off right after Proteales (*Nelumbo*). Trees based on concatenation, MRL, and ASTRAL-CSSC tree in general also support Vitales as another clade of early-diverging eudicots, but from these trees, Santalales also clustered with Vitales (Supplemental Figures 2-5, and 7). Although several previous large-scale phylogeny studies have reported the positioning of Vitales as sister to core-eudicots or early-diverging eudicot, this also has been regarded as an artefact (Wickett et al., 2014; Zeng et al., 2014). A recent synteny analysis using the columbine genome and grapevine genome revealed that the gamma palaeohexaploidy at the root of core-eudicots is likely a result of hybridization between a tetraploid and a diploid species, thus the core-eudicots have a hybrid origin (Aköz and Nordborg, 2019). This conclusion could also help us better understand the phylogenetic incongruence of the basal groups within core-eudicots including Vitales.

For monocots, the synteny tree proposed some rather unexpected relationships regarding the BEP and PACMAD clade (of Poales), and breaks the monophyly of the BEP clade (Difference #6 in Figure 2). To our knowledge, similar results have rarely been reported, however the comparative phylogenomic large-scale gene family expansion and contraction analysis from the wheat genome project showed a similar clustering pattern of species based on gene family profiles, thus provided some evidence in support of our result (Figure 4A of International Wheat Genome Sequencing Consortium, 2018). If the synteny tree is indeed true, it would reshape our understanding of the evolution of some of the most important crops including wheat, rice, and maize, as well as the origin of the C4 lineages (Sage et al., 2012). As discussed above, hybridization can lead to massive introgression that obscures species boundaries, thus it is plausible that the incongruences discussed above may again indicate an effect from introgression caused by (recent or more ancestral) hybridization. In conclusion, our study does not only provide a means to reconstruct species trees based on (micro)synteny from large volumes of available whole-genome data, but might also provide alternative interpretation and novel insights into elusive phylogenetic relationships and into our understanding of angiosperm evolution.

## Materials and Methods

### Genome resources

Reference genomes were obtained from public repositories, including Phytozome, CoGe, GigaDB, and NCBI. For each genome, we downloaded FASTA format files containing protein sequences of all predicted gene models and the genome annotation files (GFF/BED) containing the positions of all the genes. We modified all peptide sequence files and genome annotation GFF/BED files with corresponding species abbreviation identifiers. After constructing our synteny network database and clustering (see further), poor quality genomes could be relatively easy identified (see Zhao and Schranz 2019), and were removed from the database for further analysis. After quality control, the final list of genomes used in the current study and related information for each genome can be found in Supplemental Table S1.

### Synteny network construction and network clustering

The synteny network construction method has been described in detail before (Zhao and Schranz, 2019). The pipeline consists of two main steps: first, all-vs-all reciprocal annotated-protein comparisons of the whole genome using DIAMOND was performed (Buchfink et al., 2015), followed by MCScanX (Wang et al., 2012), which was used for pairwise synteny block detection. Parameter settings for MCScanX have been tested and compared before (Zhao and Schranz, 2019); here we adopt ‘b5s5m25’ (b: number of top homologous pairs, s: number of minimum matched syntenic anchors, m: number of max gene gaps) for flowering plant genomes. Further details regarding phylogenomic synteny network construction can be found in the Github tutorial (https://github.com/zhaotao1987/SynNet-Pipeline). Each pairwise synteny block represents pairs of connected nodes (syntenic genes), all identified synteny blocks contain millions and millions of edges, which form a comprehensive synteny network with millions of nodes. In this synteny network, nodes are genes (from the synteny blocks), while edges connect ‘syntenic’ genes.

The entire synteny network (database) needs to be clustered for further analysis. We used the Infomap algorithm for detecting synteny clusters within the map equation framework (Rosvall and Bergstrom, 2008) (https://github.com/mapequation/infomap). We used the two-level partitioning mode with ten trials. The network was treated as undirected and unweighted. Resulting synteny clusters vary in size and composition. A typical synteny cluster comprises of syntenic genes shared by groups of species, which precisely represent phylogenetic relatedness of genomic architecture among species (Figure 1b).

### Phylogenomic profiling, building the synteny matrix, and maximum-likelihood estimation (Syn-MRL)

A cluster phylogenomic profile shows its composition by number of nodes in each species. We summarize the total information residing in all synteny clusters as a data matrix for tree inference. Phylogenomic profiles of all clusters construct a large data matrix, where rows represent species, and columns as clusters (Figure 1b). The matrix was then binarized (i.e. #nodes ≥ 1 was defined as 1, #nodes = 0 was defined as 0) as the final synteny matrix (Figure 1b), and converted into ‘phylip’ format for phylogenetic inference.

Tree estimation was based on maximum-likelihood as implemented in IQ-TREE (version 1.7-beta7) (Nguyen et al., 2014), using the MK+R+FO model. (where “M” stands for “Markov” and “k” refers to the number of states observed, in our case, k = 2)

The +R (FreeRate) model was used to account for site-heterogeneity, and typically fits data better than the Gamma model for large data sets (Yang, 1995; Soubrier et al., 2012). State frequencies were optimized by maximum-likelihood (by using ‘+FO’). We generated 1000 bootstrap replicates for the SH-like approximate likelihood ratio test (SH-aLRT), and 1000 ultrafast bootstrap (UFBoot) replicates (-alrt 1000 -bb 1000) (Hoang et al., 2017).

### Gene marker identification, phylogenetic tree construction, MRL, topology test

For sequence alignment-based approaches, we identified BUSCO and CSSC marker genes from genome annotated protein sequences and synteny clusters, respectively. Candidate BUSCO genes were identified using BUSCO v3.0 (embryophyta_odb9, 1440 BUSCOs) (Simão et al., 2015). The criterion for charactering genes of CSSC (conserved single-copy synteny clusters) was: median number of nodes across species < 2, present in ≥ 90% genomes, and presence within Poaceae, monocots (except Poaceae), Asterids, Rosales, Brassicaceae, and Fabaceae must ≥ 50%.

Multiple sequence alignments were performed using MAFFT (version 7.187) (Katoh and Standley, 2013). Two rounds of alignment trimming and filtering were conducted by trimAl (Capella-Gutiérrez et al., 2009). First, the alignments were trimmed through heuristic selection of the automatic method (-automated1). Second, sequences with less than 50% residues that pass the minimum residue overlap threshold (overlap score: 0.5) were removed (-resoverlap 0.5 -seqoverlap 50). Alignment concatenation was conducted by catfasta2phyml (https://github.com/nylander/catfasta2phyml).

Maximum-likelihood analyses were conducted using IQ-TREE (Nguyen et al., 2014). For sequence-based tree construction, we used the JTT+R model for protein alignments, both for the construction of trees based on single alignments, as well as for the concatenated sequence alignments (Figure 1a right panel). For all trees, we performed bootstrap analysis with 1000 bootstrap replicates (SH-aLRT and UFBoot (-alrt 1000 -bb 1000)).

ASTRAL-Pro (Zhang et al., 2019) was used for the tree summary approach based on multi-species coalescence to estimate the species trees from 1438 BUSCO gene trees and 883 CSSC gene trees, respectively. ASTRAL-Pro is the latest update of ASTRAL, which can now account for multi-copy trees.

For the MRL supertree analysis, 1438 BUSCO gene trees and 883 CSSC gene trees were used as two independent data sets (Figure 1c). For each tree of the two sets, branch support from bootstrap analysis has been included as a standard output of IQ-TREE (suffixed as ‘*.phy.splits.nex’). We then encoded all splits (bipartitions) with ≥ 85% UFBoot support of all the trees into the data matrix. Coding is similar to the “Baum-Ragan” coding method (0, 1, ?) (Baum, 1992), but without question marks because ‘?’ was originally designed for missing taxa as trees were multi-sourced. We used the same binary model (MK+R+FO) in IQ-TREE, and parameter settings were as the one described earlier in Syn-MRL supercluster analysis.

To assess whether alternative sister-group relationships of certain plant clades could be statistically rejected by synteny matrix, we performed approximate unbiased (AU) tests (Shimodaira, 2002), as implemented in IQ-TREE (Nguyen et al., 2014), under the ‘MK+R+FO’ model, with 10,000 replicates.

## Acknowledgements

Z. L. is supported by a postdoctoral fellowship from the Special Research Fund of Ghent University (BOFPDO2018001701). A. Z. is supported by a PhD fellowship of the Research Foundation Flanders (FWO).

## Supplemental documents

Supplemental Figure S1 Synteny tree

Supplemental Figure S2 Concatenation-BUSCO tree

Supplemental Figure S3 Concatenation-CSSC tree

Supplemental Figure S4 MRL-BUSCO tree

Supplemental Figure S5 MRL-CSSC tree

Supplemental Figure S6 ASTRAL-BUSCO tree

Supplemental Figure S7 ASTRAL-CSSC tree

Supplemental Figure S8 Tree based on synteny matrix without magnoliids signals

Supplemental Table S1 List of genomes used in this study

Supplemental Table S2 Phylogenomic profiling rearranged by *Cinnamomum micranthum* chromosomes

## Data availability

Datasets used in this study are available at DataVerse (https://doi.org/10.7910/DVN/7ZZWIH). This includes all annotated protein sequences in FASTA format of each genome, entire synteny network database (edgelist), network clustering result, trimmed alignments and corresponding phylogenetic trees of BUSCO and CSSC genes, bipartitions with support values for each tree, and binary data matrices. Related codes and software parameters are available at Github (https://github.com/zhaotao1987/Syn-MRL).

## Author contributions

Y.V.d.P., M.E.S, and J.X. conceived the idea. T.Z. and Y.V.d.P. designed the study. T.Z. and S.K. performed the analysis. T.Z., J.X., Y.V.d.P, Z.L., M.E.S., and A.Z. analyzed data. T.Z. and Y.V.d.P. wrote the manuscript. All authors discussed the results and commented on the manuscript.

